# Fragmented habitat compensates for the adverse effects of genetic bottleneck

**DOI:** 10.1101/2022.04.05.487133

**Authors:** Ari Löytynoja, Pasi Rastas, Mia Valtonen, Juhana Kammonen, Liisa Holm, Morten Tange Olsen, Lars Paulin, Jukka Jernvall, Petri Auvinen

**Affiliations:** Institute of Biotechnology, HiLIFE, University of Helsinki, Finland; Organismal and Evolutionary Biology Research Program, Faculty of Biosciences, University of Helsinki, Finland; Department of Geosciences and Geography, Faculty of Science, University of Helsinki, Finland; Natural Resources Institute Finland (LUKE), Helsinki, Finland; Section for Molecular Ecology and Evolution, Globe Institute, University of Copenhagen, Denmark

## Abstract

In the face of human-caused biodiversity crisis, understanding the theoretical basis of conservation efforts of endangered species and populations has become increasingly important. According to population genetics theory, population subdivision helps organisms retain genetic diversity, crucial for adaptation in a changing environment. Habitat shape is thought to be important for generating and maintaining population subdivision, but empirical cases are needed to test this assumption. We studied Saimaa ringed seals, landlocked in a labyrinthine lake and recovering from a drastic bottleneck, by whole-genome sequencing 105 individuals and additional individuals from three other ringed seal subspecies. We analyzed the distribution of variation and genetic relatedness among the individuals in relation to the habitat shape. Despite a severe history of a genetic bottleneck with prevalent homozygosity in Saimaa ringed seals, we found evidence for population structure mirroring the subregions of the lake. Highlighting the significance of habitat connectivity in conservation biology and the power of genomic tools in understanding its impact, genome-wide analyses showed that the subpopulations had retained unique variation and largely complementary patterns of homozygosity. Integration of genetic analyses in conservation decisions gives hope to Saimaa ringed seals and other endangered species in fragmented habitats.

## Introduction

Human activities have had profound impacts on the diversity of terrestrial and marine organisms in the form of habitat fragmentation, overexploitation and disturbance. Conservation efforts aim at retaining and increasing the census size (*N*) of endangered species and populations, but the significance of *N_e_*, the effective population size, is also widely acknowledged (1–3), e.g., when designing protected areas or deciding to translocate animals. The concept of *N_e_* was developed by Wright (4) and Fisher (5) and refers to the size of an idealized population that behaves genetically similarly to the real-life population under study. In conservation biology, the significance of *N_e_* is that it determines the populations’ ability to respond to evolutionary forces, e.g., to adapt to a changing environment or to survive under negative selection (6). While *N_e_* can be affected by sex ratio, age structure and differences in reproductive success, one of its key determinants is the population structure. Wright (4; 7) showed that the metapopulation effective size, *N_eT_*, is a function of *N_T_*, the total population size, and *F_ST_*, a differentiation between subpopulations, 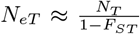. Although this suggests that the effective population size increases in structured populations, Wright made strong underlying assumptions about the nature of subpopulations, and in realistic scenarios, population subdivision is thought to reduce rather than increase the global effective size (6). Many results of Wright and Fisher’s early work have later been derived from coalescence theory (8). There, *N_e_* is defined as the coalescence time of sequences and is always increased by population subdivision.

Lake Saimaa, Finland, provides a unique opportunity to empirically study the effects of population substructure on the preservation of genetic diversity and to learn about the significance of the environment shape for the conservation efforts of endangered species in general. The Saimaa ringed seals (*Pusa hispida saimensis*) are a land-locked population of ringed seals, and along with the Arctic (*P. h. hispida*), Okhotsk (*P. h. ochotensis*), Baltic (*P. h. botnica*), and Ladoga (*P. h. ladogensis*) ringed seals, one of the five recognized subspecies of the ringed seals. According to estimates (9), the census sizes of Okhotsk and Arctic ringed seals are counted in hundreds of thousands and millions, while the census size of the Baltic population is around 10,000 and that of the Ladoga population 3,000-5,000 individuals. While the population size of Saimaa ringed seals has recovered from 100-160 individuals in the early 1980s to more than 400 individuals in 2020 (10), the consequences of the extreme bottleneck to the population’s adaptive potential and long-term prospects are unclear. Lake Saimaa is highly labyrinthine with a shore length of 13,700 km for an area of 4,400 km^2^, and is further fragmented by ca. 14,000 islands. This is in stark contrast to neighboring Lake Ladoga, housing the only other freshwater subspecies of ringed seals, with a shore length 1,570 km for an area of 17,700 km^2^. Ringed seals from different parts of Lake Saimaa have been shown to be genetically differentiated (11; 12), but the magnitude of this population stratification and its impact on the amount of genetic diversity and the subspecies’ adaptive potential has not been assessed in detail.

## Results

Since 1980s, tissue samples of deceased Saimaa ringed seals have been collected for research needs. 19 of these were whole-genome sequenced at medium (mean 11.2X) and 86 at low sequencing coverage (mean 5.2X; Table S1). To place them into a global context, 9-19 samples were sequenced from the Ladoga, Baltic, and Arctic subspecies (mean 8.3X; Fig. 1A; Table 1; Table S1). We used the 2.350 Gbp contig assembly for the Saimaa ringed seal and in most analyses focused on 55.9% of the genome defined as callable (13) and unique (14); a subset of analyses were performed on the 100 longest contigs, the shortest of these being 5.48 Mbp and them in total covering 804 Mbp (34%) of the genome. Using bwa mem (15) and samtools (16) for read mapping, ANGSD (17) and beagle (18) for genotyping and phasing, we separately processed the Saimaa samples (‘Saimaa data’; n=105) and the medium coverage samples of the four subspecies (‘subspecies data’; n=59). Among the 1.313 Mbp sites, we found in total 5,894,190 single nucleotide polymorphisms (SNPs) in the Saimaa data and 9,830,328 SNPs in the subspecies data.

**Table 1:** Sample size, sequencing coverage, and number of variants for the four ringed seal subspecies.

|  | Arctic | Baltic | Ladoga | Saimaa |
| --- | --- | --- | --- | --- |
| samples | 19 | 12 | 9 | 19 |
| coverage | 5.1X | 11.3X | 10.8X | 11.2X |
| total SNPs | 8,609,960 | 8,731,833 | 7,979,288 | 4,922,410 |
| private SNPs | 79,879 | 24,304 | 20,265 | 257,481 |
| private fixed | 17 | 9 | 71 | 6,495 |

**Fig. 1:**
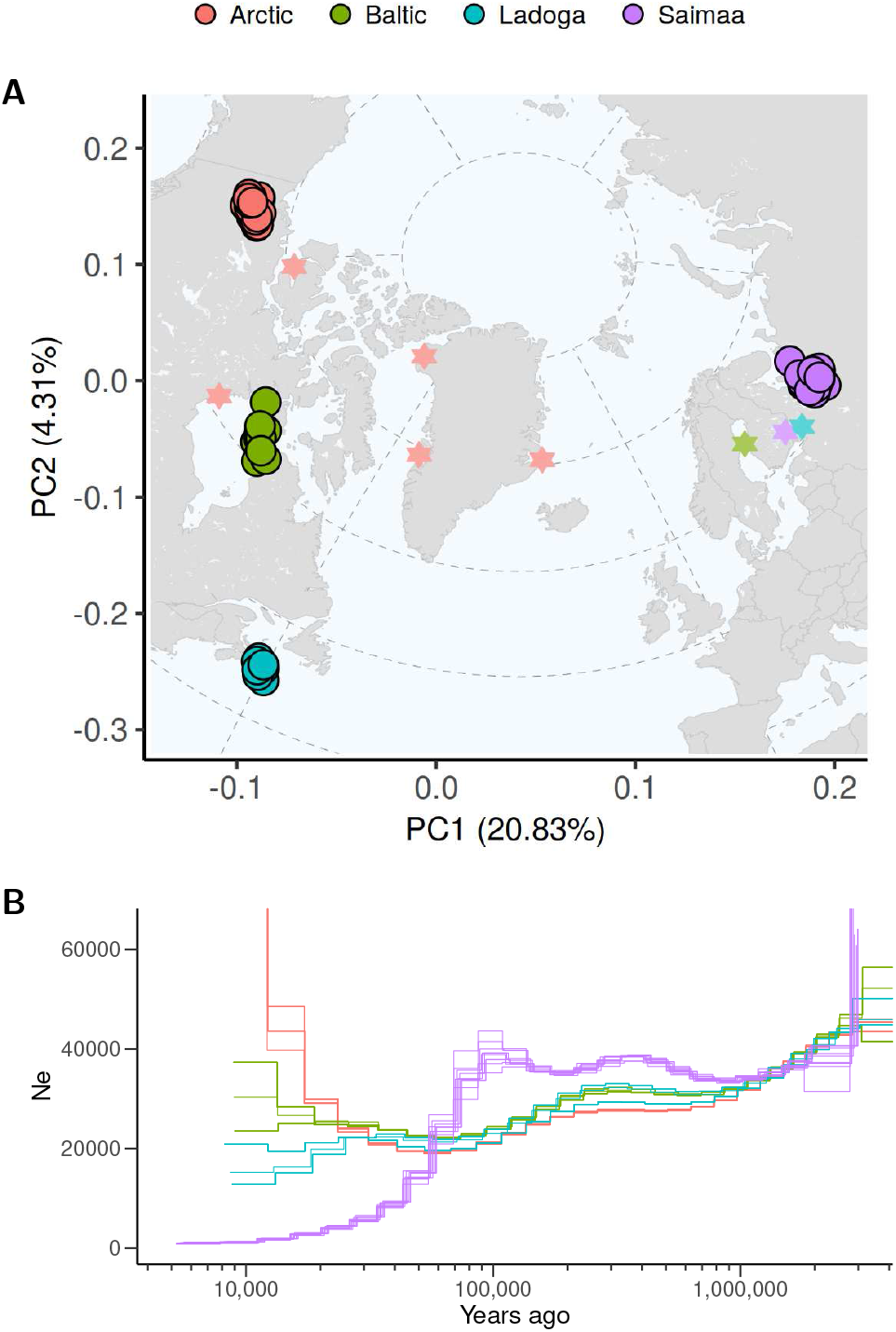
Genetic divergence and demographic histories of the four ringed seal subspecies. (A) Dot plot shows the first two axes of PCA of genetic variation (dots), and background map the origin of the samples (stars). (B) The lines indicate inferred ancestral population sizes (*N_e_*) across time. For each subspecies, the analysis was performed for ten randomly chosen sample pairs.

### Saimaa subspecies is genetically diverged from the others

In the principal component analysis (PCA) of the subspecies data, the first PC explains 20.8% of total genetic variation and separates the Saimaa from all others; the second PC explains 4.3% and separates the three other subspecies (Fig. 1A). Despite smaller sample sizes (n=9-19), the Arctic, Baltic and Ladoga subspecies contain significantly greater numbers of variants, from 7.98 × 10^6^ to 8.73 × 10^6^ SNPs each, than the Saimaa samples (n=19), in total 4.92 × 10^6^ SNPs (Table 1). 32.9% of the variants are shared between all subspecies, while an additional 37.0% is shared between the three non-Saimaa subspecies (Fig. S1). Consistent with this, the Saimaa sample contains more than 257,000 private SNPs and 6,485 fixed differences, while the other three subspecies have 20,200-69,900 private SNPs and 8-71 fixed differences (Table 1, Fig. S1).

PCA and SNP statistics indicate that the Saimaa population has diverged from other ringed seals and lost genetic diversity in comparison to other subspecies. To understand this in more detail, we inferred the demographic histories with MSMC2 (19) using the mutation rate estimated for polar bears, 1.826 × 10^−8^ substitutions per site per generation (20), generation time of 10 years (21) and ten four-chromosome sets from each subspecies. The *N_e_* trajectories for Arctic, Baltic and Ladoga subspecies are very similar until 25,000 years ago (ya; Fig. 1B); then, the Arctic population grows strongly while the Baltic population increases slightly and the Ladoga population decreases in size. The trajectories of Saimaa individuals separate from others around 1,000,000 ya and show a clearly higher *N_e_* until 100,000 ya. After that, the Saimaa *N_e_* dives and eventually drops below 1,000, more than an order of magnitude lower than for any other sub-species (Fig. 1B).

Three points are noticeable. First, the diverging estimates of recent *N_e_* for Arctic and Baltic (Fig. 1B) disagree with their very similar SNP statistics (Table 1) and suggest that, due to the low sequencing coverage, our SNP counts miss recent variants, affecting most the Arctic population. The MSMC method uses the lengths and ages of coalescent blocks (19) and is not significantly affected. Second, a growth in MSMC estimates can be explained either by an increase in the actual census size or by population structure and admixture between distinct lineages (4; 7; 6; 8), and the bump in the Saimaa *N_e_* before 100,000 ya (Fig. 1B) possibly reflects the population deriving some of its ancestry from elsewhere. Similar signals are seen in other aquatic organisms in the Baltic Sea region (22–24), and an origin in an eastern freshwater refugium has been proposed also for Saimaa ringed seals based on their unique mitochondrial haplotypes (25). Finally, conversion of coalescent estimates into years and numbers of individuals requires a good understanding of the genomic mutation rate, and lacking that, our estimates should be considered approximate. This does not invalidate the clear qualitative differences between Saimaa and three other subspecies, however.

### Saimaa seals have lost much of their genetic variation

Genetic variation can be quantified as heterozygosity of individuals (*H*), nucleotide diversity within populations (*π*) and genetic divergence between populations (*d_xy_*). Genome-wide mean estimates confirm the lower variation in the Saimaa subspecies, with *H* and *π* of 0.0012 and 0.0013 differences per site, respectively, being slightly more than half of that of the Baltic and Ladoga subspecies (Table 2). On the other hand, the highest *d_xy_* estimates are for pairs involving Saimaa, consistent with the population containing private variation. Due to the lower sequencing coverage (Table 1), the diversity estimates for the Arctic subspecies are likely downward biased (Fig. S2). The distributions of local estimates *H* and *π*, computed in 250 kbp genomic windows, peak around 0.002 differences per site for the three non-Saimaa subspecies and are nearly indistinguishable from each other (Fig. 2A,B). The picture is very different for Saimaa and, while the distribution of *π* peaks around 0.0012 (Fig. 2B), *H* is concentrated close to zero with another gently sloping bump above 0.002 (Fig. 2A). This indicates that the genomes of individual Saimaa seals contain long segments with virtually no variation, but copies of these low-variation regions tend to differ among the seals.

**Table 2:**
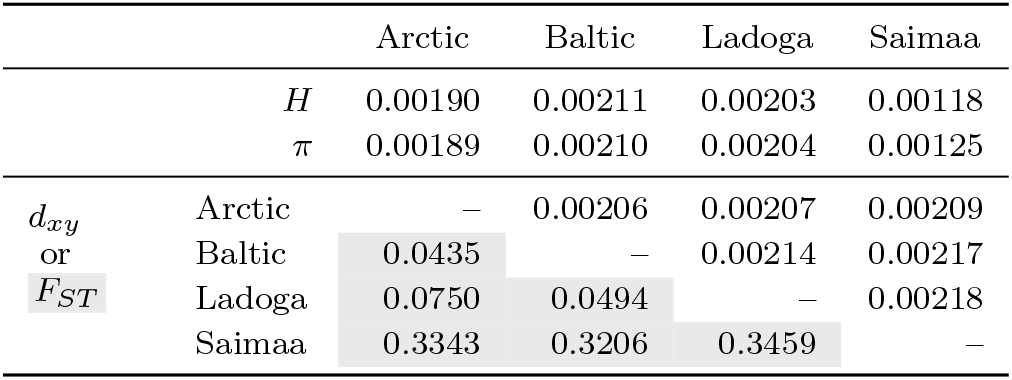
Heterozygosity of individuals (*H*), nucleotide diversity within populations (*π*), genetic divergence between populations (*d_xy_*; above diagonal) and population differentiation (*F_ST_* ; below diagonal) for the four ringed seal subspecies.

|  |  | Arctic | Baltic | Ladoga | Saimaa |
| --- | --- | --- | --- | --- | --- |
| | $H$ | 0.00190 | 0.00211 | 0.00203 | 0.00118 |
| | $\pi$ | 0.00189 | 0.00210 | 0.00204 | 0.00125 |
| $d_{xy}$<br>or<br>$F_{ST}$ | Arctic | — | 0.00206 | 0.00207 | 0.00209 |
|  | Baltic | 0.0435 | — | 0.00214 | 0.00217 |
|  | Ladoga | 0.0750 | 0.0494 | — | 0.00218 |
|  | Saimaa | 0.3343 | 0.3206 | 0.3459 | — |

**Fig. 2:**
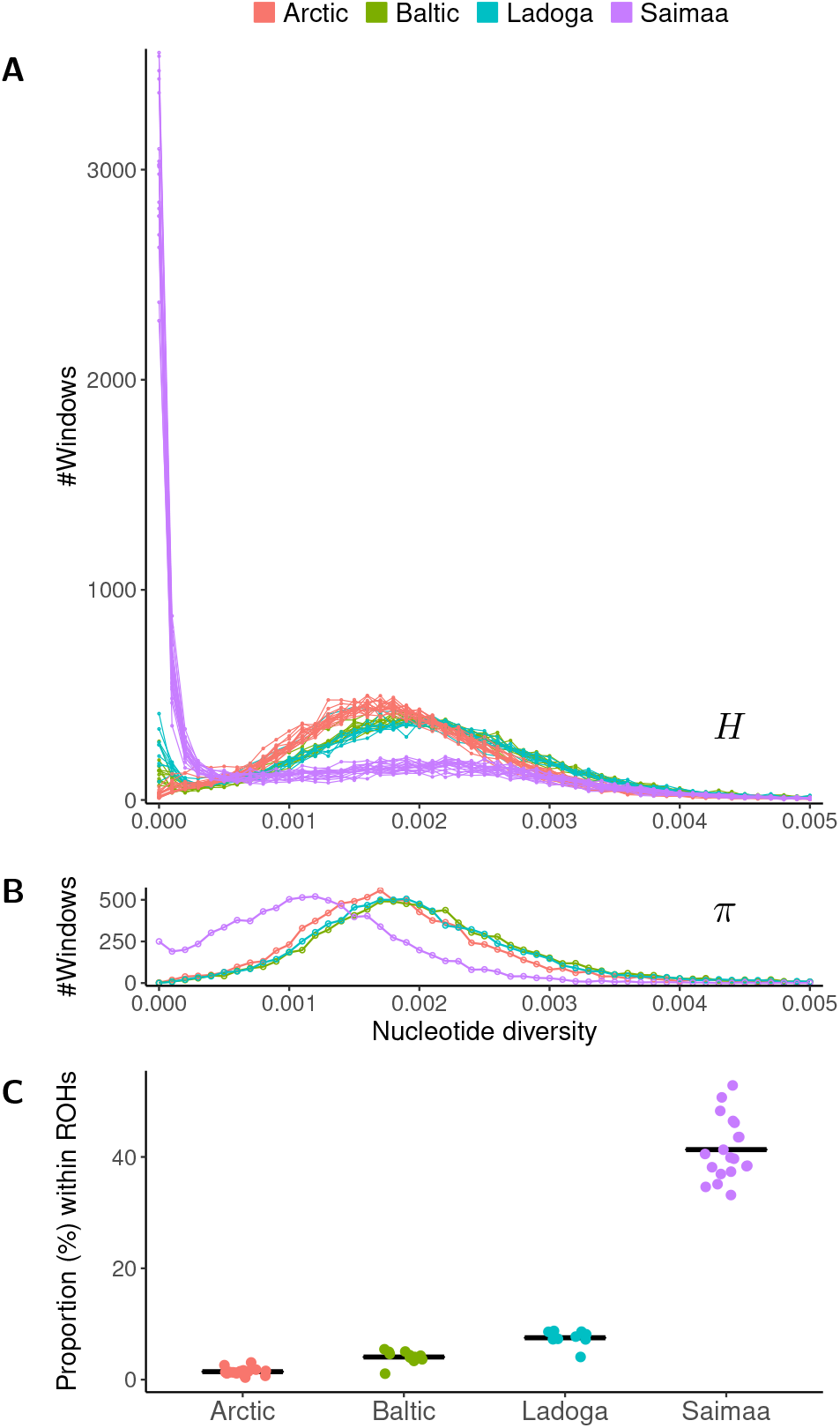
Nucleotide diversity and runs of homozygosity across the genomes. (A, B) Number of 250 kbp genomic blocks (y axis) with nucleotide diversity in given bracket (x axis) for (A) individuals (heterozygosity; *H*) and (B) populations (*π*). Different subspecies are indicated with colors. (C) Proportion of the genome (y axis) within ROHs of at least 100 kbp in length for individuals of different subspecies. Vertical lines indicate the sample means.

The lack of genetic variation implicates inbreeding and formation of runs of homozygosity (ROH) in the genome. Consistent with their largest *N_e_*, Arctic individuals have the smallest amount, on average less than 1.44%, of their genome within ROHs longer than 100 kbp (Fig. 2C). The Ladoga seals have a higher proportion of the genome covered by ROHs than the Baltic ones (7.53% vs. 4.07%), though neither show signs of contemporary inbreeding (26). In contrast to this, in Saimaa individuals between 33.2 and 52.9% of the genome are within ROHs greater than 100 kbp (Fig. 2C).

In addition to the lower absolute sequencing coverage for the Arctic samples, variant imputation (applied to compensate for the low coverage) and phasing may perform differently on populations with vastly different effective sizes. This has no impact on the big picture, though, and Saimaa clearly has a deep independent history and its size has remained small for a long time. However, the lake is extremely fragmented and previous studies have shown that seals from different parts of the lake are genetically diverged (11; 12). For the conservation efforts, it is crucial to understand the impact of the small population potentially breaking into even smaller subpopulations.

### The small Saimaa population is highly structured

The presence of substructure within Lake Saimaa was confirmed with a PCA of the 98 Saimaa individuals with known geographic information: the first two PCs explain 8.6 and 4.7% of genetic variation and separate, respectively, the individuals from Northern Saimaa (NS) and Southern Saimaa (SS) from all others, leaving Central Saimaa (CS) to connect the different clusters (Fig. 3A). With one exception, the three groups occupy distinct lake regions connected by narrow straits (Fig. 3B). Additional PCs separate individuals from specific regions such as Kolovesi, the most fragmented and labyrinthine part of the lake (Fig. S3A-E).

**Fig. 3:**
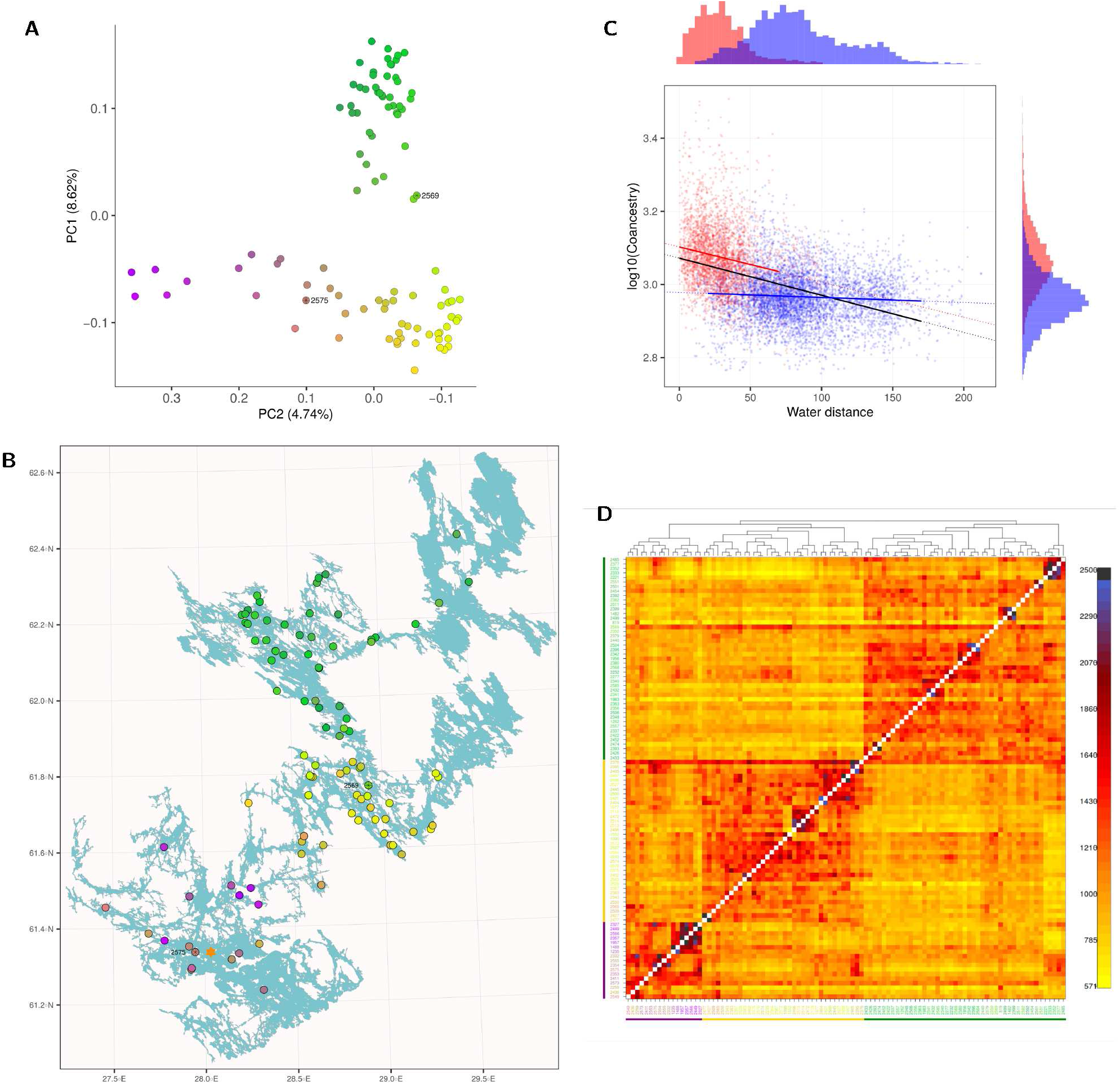
Genetic variation within Saimaa. (A) First two PCs explain 8.6 and 4.7% of the variation among the 98 samples. (B) The map of Saimaa shows the fragmentation and the labyrinthine shape of the lake. The sampling locations of the studied seals are indicated with dots matching the colors in the PCA plot. Orange star indicates the living area of a seal translocated in 1992 and black asterisks mark individuals ‘2569’ and ‘2575’. (C) Logl0-transformed coancestry estimates and water distance between sample pairs correlate but this is largely explained by pairs coming either from the same or from different lake regions, indicated by red and blue points, respectively. Histograms on top and right show the distribution within and between lake regions. The black, red and blue regression lines are fitted to full data, to red dots and to blue dots, respectively. (D) The coancestry matrix contains three blocks of higher similarity (scale on the right with higher values indicating greater genetic similarity) that, with the exception of ‘2569’, consist of samples from northern (top, indicated by green line), central (middle, yellow) and southern (bottom, magenta) part of the lake. The dendrograrn indicates the inferred grouping of the samples, and the axis label colors (left, bottom) match those in the PCA plot.

To test whether the distribution of variation is explained by simple isolation by distance, we computed water distances between the map locations and the genetic similarities between the samples, using coancestry as defined by fineSTRUCTURE (27). The logarithmic transformation of coancestry (27) correlates strongly with geographic distances (r=−0.441, *p* < 2.2 × 10^−16^). However, analysis of partitioned data reveals that this correlation is driven by both genetic similarities and physical distances being very different within and between different lake regions (Fig. 3C): within regions, the correlation is strong (cor=−0.200, *p* < 2.2 × 10^−16^), but for sample pairs coming from different lake regions, it is almost non-existent (cor =−0.070, *p* < 4.7 × 10^−8^). Consistent with this, the coancestry matrix contains three large blocks of high relatedness, the corresponding individuals coming from the three distinct parts of the lake, but also blocks of a few individuals with very high relatedness (Fig. 3D). In addition to blocks, the matrix contains stripes indicating genetic identity to many other seals. One of these individuals, ‘2569’, is the clearest deviation among the three groups and is firmly amongst the Northern samples in the PCA (Fig. 3A) despite being originally sampled deep within the Central region of the lake (Fig. 3B). ‘2569’ shows coancestry to different lake regions (Fig. S4A) and this mixed ancestry produces one of the highest levels of genomic heterozygosity (Table S2).

### Saimaa subpopulations contain private variation

To quantify the current variation within subpopulations, we extracted the 22 Saimaa samples born in the 2010s (Fig. S5). Sample ‘2569’ was excluded as an apparent migrant and two others due to atypically high homozygosity, possibly contributing to their deaths as young pups. Consistent with the coancestry estimates, Northern samples contain more genetic variation than Central and Southern samples, and each subpopulation contains some private variation, as indicated by *d_xy_* being higher than *π* (Table 3). The *F_ST_* estimates, measuring population differentiation, between the subpopulations vary from 0.086-0.114 (Table 3), and are thus of the same magnitude as between the human superpopulations (28). *F_ST_* analysis is known to depend heavily on SNP ascertainment and estimates from different data sets are not fully comparable (28). Nevertheless, in our analyses the Saimaa subpopulations (Table 3) show greater population differentiation than the three other ringed seal subspecies (Table 2). The *F_ST_* estimates are not markedly different with the full 98 sample set, varying from 0.082-0.106.

**Table 3:**
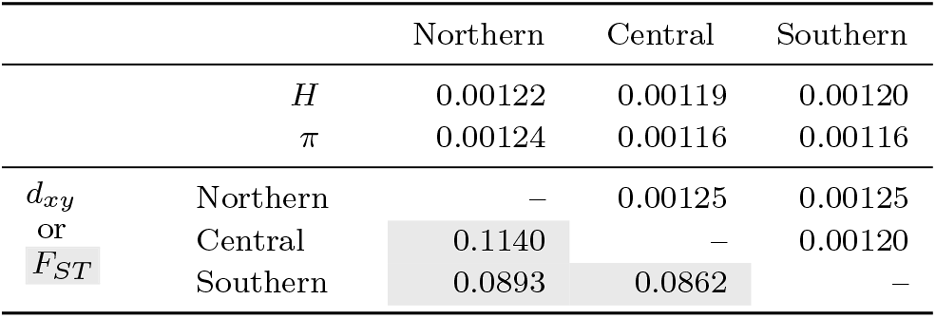
Heterozygosity of individuals (*H*), nucleotide diversity within populations (*π*), genetic divergence between populations (*d_xy_*; above diagonal) and population differentiation (*F_ST_* ; below diagonal) across the whole genome for the three Saimaa ringed seal subpopulations. Only the 22 samples born in the 2010s were included.

In analyses of 250kbp windows, the distributions resemble those of the four subspecies analysis, and H concentrates close to zero (Fig. 4A) and *π* peaks around 0.0012 (Fig. 4B). The peak around zero for *π*, meaning complete loss of variation within a window, is the greatest in Southern individuals, reflecting the low historical and current population size in that part of the lake. Crucially, *π* for the whole Saimaa shows significantly fewer genomic blocks void of any variation, a drop of 44.2% for windows with *π* below 0.0001 in comparison to the subregion mean, and in average 10.4% increase for windows with *π* between 0.0009 and 0.0015 (Fig. 4B,C). This shows that the Saimaa subpopulations have lost variation in different parts of the genome and contain private variation not present in other subpopulations.

**Fig. 4:**
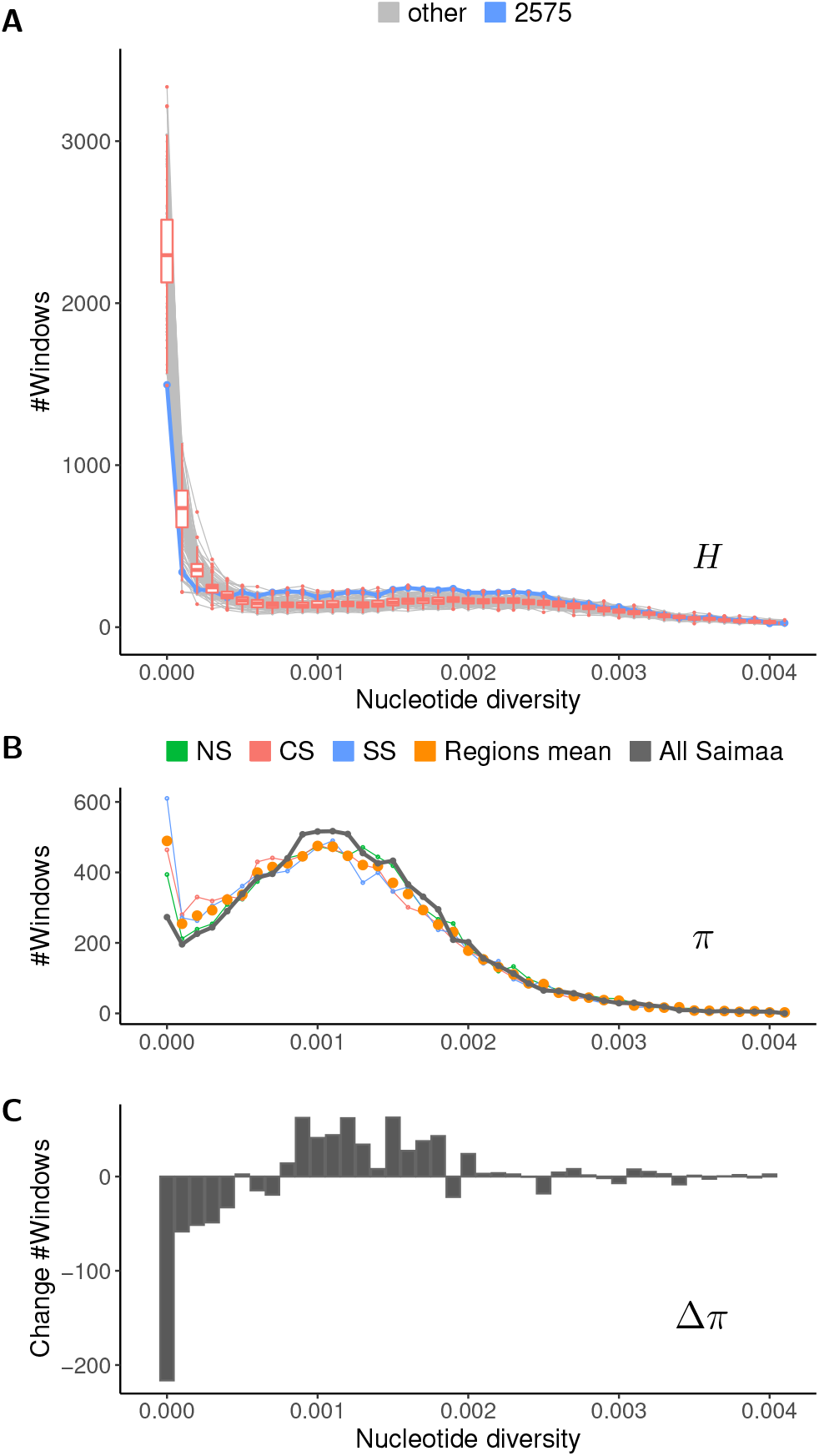
Nucleotide diversity within 250 kbp genomic blocks in Saimaa. (A) Number of genomic blocks (y axis) with heterozygosity in given bracket (x axis) for ‘2575’ (blue) and all other Saimaa individuals (light gray) and the variation of the latter (red boxplots). (B) Number of genomic blocks with *π* in given bracket for all Saimaa (dark gray) and the average (orange) of different Saimaa regions (NS, CS, SS). (C) Change in number of windows falling in each diversity bracket in the total population and the three subpopulations, i.e., the difference between the gray and orange lines.

We evaluated the significance of population substructuring using synthetic data reflecting the properties of the Saimaa data (Fig. S6). Overall, whereas increasing number of subpopulations with limited migration leads to the loss of genetic variation in individual subpopulations, the total amount of variation retained in the whole population actually increases (Fig. S7). Interestingly, three distinct subpopulations, indicating a relatively low migration rate, observed in Lake Saimaa seem to be close to the sweet spot where the subpopulations do not accumulate too many ROHs while the total population retains more variation than one panmictic population (Fig. S7). With only two subpopulations and high migration, the benefits of substructuring over a panmictic population are negligible; with greater numbers of subpopulations or very low migration rates, the high homozygosity starts to threaten the survival of the subpopulations (Fig. S7).

### Coancestry reveals a past translocation experiment?

One case of migration within Saimaa is empirically known as an adult female was translocated from the north-western part to the southern part of the lake in 1992 (29). No tissue sample is available, but the female is known to have reproduced in the area since 1994 (29) and was still alive in 2020 (10). Interestingly, the PCA shows the seals around her living area to be genetically intermediate while the most distinctively “southern” seals are from the narrow entry through which the possible natural migration has to enter that part of the lake (Fig. 3A,B). There are 73 sample pairs with coancestry above 2000, all within the distinct parts of the lake (Fig. 3D, Fig. S4B). On the other hand, there are 29 cross-region sample pairs with coancestry above 1500 (Fig. S4C) and these include ‘2575’ from the immediate vicinity of the translocated seal’s living area (Fig. 3B), showing the highest genetic similarity from the southern part to the northern part of the lake (Fig. S4D). A closer look at the distribution of *H* reveals that ‘2575’ has a notably lower proportion of genomic segments with no variation (Fig. 4A), consistent with the individual having mixed ancestry and containing genetic variation from different parts of the lake.

## Discussion

Lake Saimaa is unique in its fragmentation and labyrinthine outline. For a comparable area, the lake shoreline is roughly 58 times longer than if the lake would be a perfect circle. The labyrinthine shape of Lake Saimaa, together with its isolation, provides a natural setting to examine how population substructure responds to habitat fragmentation. Especially considering large mammals that typically require broad home ranges, Saimaa ringed seals offer a test case to measure the genetic effects of population subdivision.

Our analyses show that Saimaa ringed seals are unique among the studied ringed seal subspecies and contain a high number of private polymorphic sites and fixed nucleotide changes, consistent with the proposed long isolation and population bottleneck in Saimaa (25). Our demographic analyses reveal the severity of this bottleneck, give an approximate timing for the start of isolation and suggest a complex ancestry for the Saimaa population. Although the impact of the population bottleneck is evident in the total lack of variation in parts of the Saimaa genomes, a larger sample reveals a division into three strongly differentiated subpopulations. Importantly, individuals and subpopulations have lost variation across different genomic regions and, as a total, the Saimaa population retains some variation throughout most of the genome.

Detailed analyses show that subpopulation divisions can be linked to narrow straits that decrease migration. In the analysis of genetic variation, the first principal component separates the northern subpopulation from the rest, the split coinciding with the narrow straits surrounded by the city of Savonlinna. The second principal component separates the southern subpopulation, leaving the central part to connect the two extremities; other components separate geographically isolated individuals (cf. 12). While the subpopulation structure is suggestive of a star-like formation where peripheric parts of the lake are connected through a central population, the subpopulation in central Saimaa does not show the greater genetic variation as suggested by the simulation, possibly reflecting a lower historical population size.

The loss of genetic variation in Lake Saimaa seems drastic, but the situation could be even worse. Our simulations examining different degrees of migration between subpopulations suggest that the Saimaa population is close to the sweet spot where the population as a whole is retaining or even gaining variation due to substructuring. Although the unique shape of Lake Saimaa and its impact on population structure and migration patterns are natural, one can draw parallels to extensive work on ecosystems and the importance of the number and connectivity of habitat patches (e.g. 30). We emphasise that our study should not be interpreted as an endorsement of artificial division of large habitats but rather as a further example of the importance of migration and the value of even small subpopulations. Although Saimaa ringed seals are a very special case of habitat fragmentation with some migration, our findings should have both theoretical and practical significance in conservation genetics. Specifically, compared to previous reports on the substructuring of the Saimaa population (12) and population analyses in general (31), our analyses also demonstrate that simple metrics such as *F_ST_* alone provide limited information. Deciphering the genetic details underlying differentiation among subpopulations and individuals can reveal crucial knowledge for a species’ conservation.

Saimaa-ringed seals appear to be a success story of conservation biology and the population size of the subspecies once listed as “Critically Endangered” (32) has grown from 100-160 individuals in the early 1980s to over 400 in 2020 (10). Despite recent growth, the population remains highly vulnerable and especially warmer winters and shorter ice and snow cover pose a serious risk to the breeding and early life of the pups (10). Nevertheless, our analyses indicate that the labyrinthine shape of Lake Saimaa has benefitted the retention of genetic diversity of the population as a whole, a result that can be useful in population management strategies that include translocations.

## Methods

### Draft reference genome

The Saimaa reference individual was sequenced with Pacific Biosciences RSII SMRT sequencing to 50X coverage, and assembled into 2,183 contigs using FALCON (v.0.3.0; 33). The contig assembly was further error corrected by mapping 100X coverage of Illumina short reads into the assembly and running the Pilon (v.1.16) improvement pipeline (34). A detailed description of the Saimaa ringed seal reference genome will be given by Kammonen et al. (in preparation).

### Callable regions

The callable regions were defined with SNPable (13) and RepeatMasker (14). The SNPable mask file was generated using bwa (v.0.7.16; 15), k-mers of 35 bp and threshold of 50%. The fasta-formatted mask was converted to a bed-formatted positive mask using makeMappabilityMask.py from the MSMC tools (19). The positive mask covered 86.7% of the genome. RepeatMasker (v.open-4.0.5) was run using the rmblastn (v.2.6.0 +) search engine and RepBase Update 20150807, and query species “Canis lupus familiaris”. RepeatMasker masked 42.50% of the genome as repeats. After combining the two masks with bedtools, 55.9% of the genome was defined as callable and unique.

### Population sequencing

DNA sequencing of Baltic, Ladoga and Saimaa samples was performed on NextSeq 500 and NovaSeq 6000 Illumina platforms at the DNA Sequencing and Genomics Laboratory, Institute of Biotechnology, University of Helsinki. Arctic samples were sequenced on Illumina 4000 platform at the GeoGenetics Sequencing Core, Globe Institute, University of Copenhagen.

### Variant calling

Bam files were constructed with bwa mem followed by GATK IndelRealigner (v.3.8-0; 35). Variant calling was done with ANGSD (v.0.930; GATK genotyping mode: -GL 2; 17) and PCANGSD (v.0.981; number of PCA axes: -e 2 for the Saimaa dataset, -e 4 for the four subspecies dataset; 17) following the program documentation. The output of PCANGSD was converted to a vcf file using a custom script. Beagle (v.4.1; 36) was run on the vcf file in its low coverage mode (gl=input.vcf.gz) and then in its genotype mode (gt=beagle output.vcf.gz); for the Saimaa data, Beagle was run on all samples and for each subpopulation separately. The subsets of sites were extracted with bcftools (v.1.9-87; 16).

### Runs of homozygosity

ROHs were called with bcftools roh (v.1.9-87; 37) using the complete data and assuming a PL of 30 for the two least likely genotypes. The coverage and overlap of ROH segments of different length was computed with R package GenomicRanges (38). For the sum of ROHs, only contigs with length over 100 kbp were included, covering 98.9% of the total length of the assembly.

### Nucleotide diversity and *F_ST_*

Genome-wide heterozygosity per sample and genome-wide nucleotide diversity for the full population was computed using bcftools and custom R code. Estimates were corrected for callable sites within windows of 250,000 bp, retaining windows with at least 40% of sites inferred as callable. Genome-wide mean heterozygosity and mean nucleotide diversity were computed across all callable sites. *F_ST_* was computed with vcftools (v.0.1.13; 39) across all callable sites.

### Coancestry, IBS distances and principal component analysis

Coancestry (chromosome sharing) was computed with fineSTRUCTURE (v.4.1.1; 27) using the full data for the 100 longest contigs. The principal component analysis (PCA) was performed with smartpca (v.16000) from the EIGENSOFT package (40) using the 100 longest contigs. The geographic data for the background map was from the rnaturalearth R package (https://github.com/ropensci/rnaturalearth).

### Water distances

The lake Saimaa map was downloaded from the Finnish Environment Institute open data service (http://paikkatieto.ymparisto.fi/lapio/latauspalvelu.html) and the locations of the seal carcasses were received from the Natural Resources Institute Finland. The Saimaa map was converted to 576×717 raster (0.25km spacing), using R packages raster (https://rspatial.org/raster) and fasterize (https://github.com/ecohealthalliance/fasterize), and the shortest paths between the carcass locations were computed with R package gdistance (https://github.com/DistanceDevelopment/Distance), allowing eight movement directions.

### Data simulation

Data simulations were performed with msprime (v.0.7.4; 41). Simulated populations started with 10,000 individuals and were reduced to 1,000 individuals 1600 generations ago; after 1200 generations the populations were split into N (*N* =1-4) subpopulations and, during the last 400 generations, reduced to 300 individuals; in the end, 120 individuals were sampled. (In this setup, one “subpopulation” represents the panmictic scenario.) We assumed mutation and recombination rates of 5 × 10^−8^ and 1 × 10^−8^ events per site per generation, migration rate (i.e. the fraction of population migrating on each generation) of 0.001, 0.005, 0.01 and 0.05, and simulated 25 replicates, each consisting of 330 pieces of 3.75 Mbp long sequence (in empirical data 50% sequence is in contigs longer than 3.715 Mbp). This gave a total length of 1237.5 Mbp (empirical 1231.5 Mbp), and in total 8,250 pieces of 150 kbp genomic blocks (in empirical data, 8,478 blocks and in average 145,260 sites). From each simulation outcome, we computed *π* within 150 kbp blocks for the subpopulations and for the total population, and compared the distributions of counts of genomic windows falling within each bracket of nucleotide diversity, reflecting the analyses in Fig. 4.

## Data and code availability

DNA sequencing data and draft contig assembly have been submitted to ENA under accessions PRJEB51012 and PRJEB51013, respectively. The code for msprime simulation and computation of nucleotide diversity estimates is available at https://github.com/ariloytynoja/saimaa-population-structure.

## Acknowledgements

We acknowledge University of Eastern Finland and Metsähallitus for the Saimaa samples, M. Verevkin for the Ladoga samples, M. Kunnasranta, P. Timonen, S. Oksanen, and J. Vierimaa for the Baltic samples, and A. Rosing-Asvid, S. Ferguson and R. Dietz for the Arctic samples; the DNA sequencing and genomics laboratory for DNA isolation and sequencing; CSC – IT Center for Science Ltd. for computational resources; and Mervi Kunnasranta, Claudius Kratochwil and Atte Moilanen for discussions and comments. JJ and PA are funded by the Jane and Aatos Erkko Foundation (4-2013, 5-2017) and PA by the LIFE programme (LIFE19NAT/FI/000832).

## Authors contributions

AL, PA and JJ conceived the paper. AL designed and performed the analyses, and wrote the first draft. PR imputed and genotyped the data. MV advised in analyses of Saimaa population structure. JK assembled the reference genome. LH contributed to genome annotation. MOT provided the Arctic data. LP and PA coordinated DNA sequencing. AL and JJ finalized the text. JJ and PA head the Saimaa ringed seal genome project. All authors commented on the manuscript.

---

**Fig. S1:**
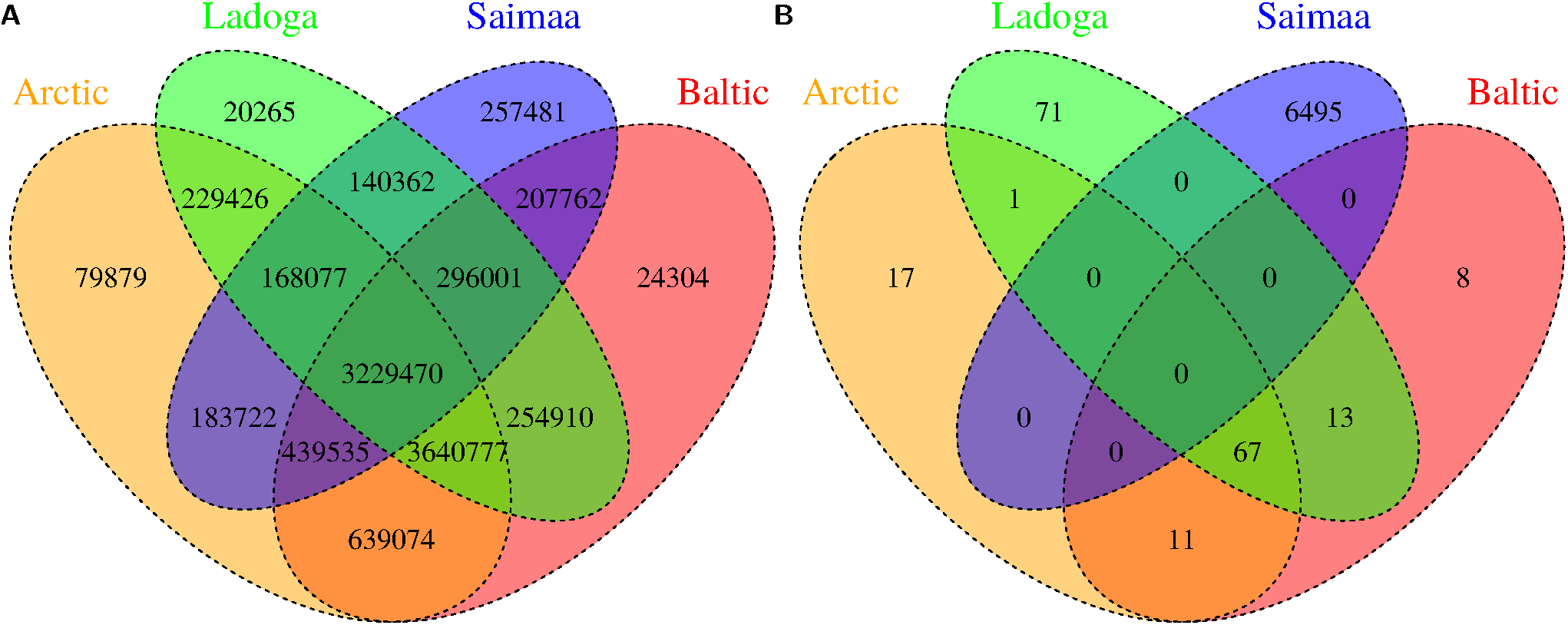
SNP sharing among the four subspecies. (A) Sharing of the 10,123,096 polymorphic sites. (B) Sharing of the 9,059 fixed sites. Together the two sets make the 10,132,155 variable sites after MAF 2.5% filtering.

**Fig. S2:**
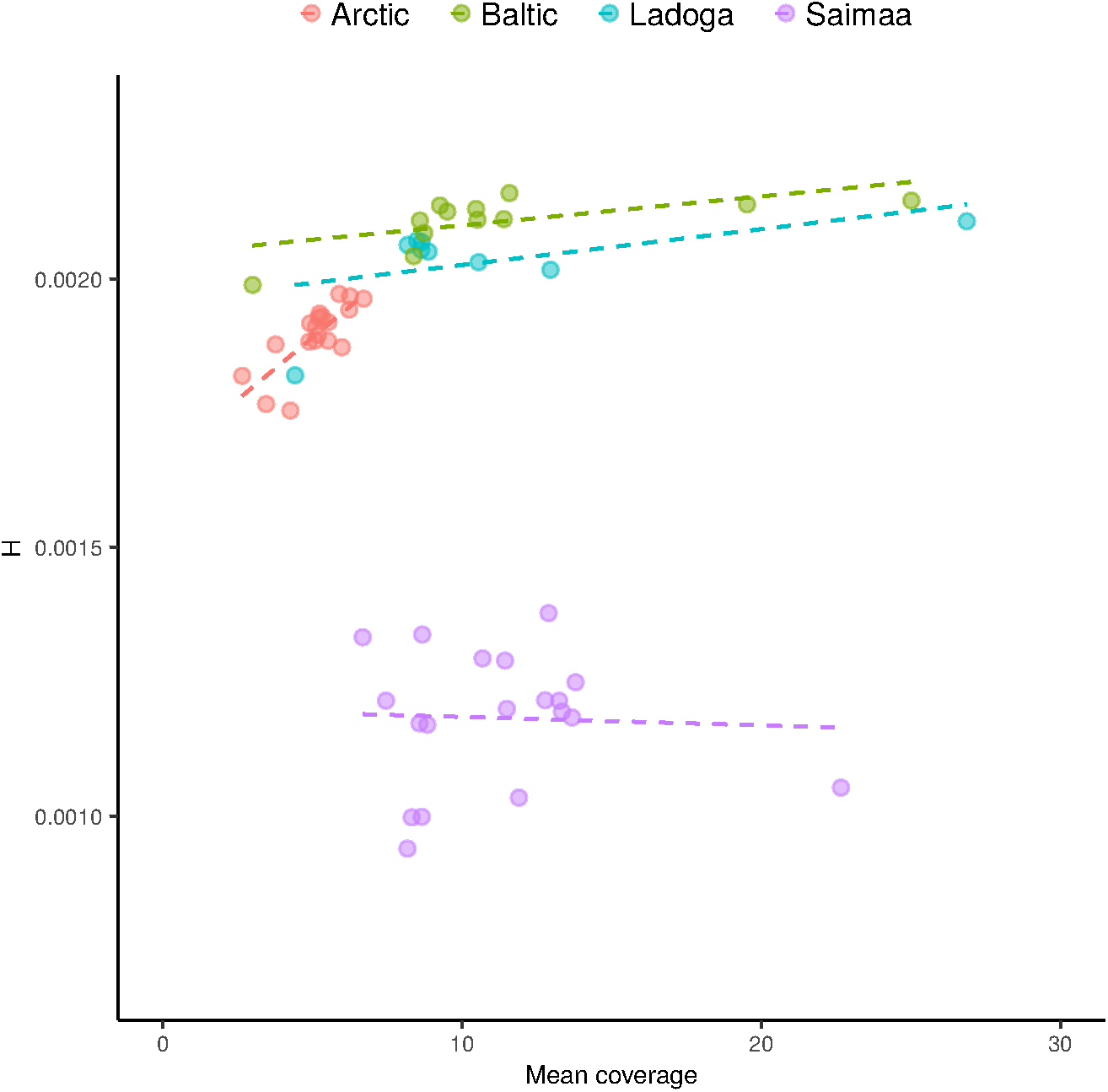
Low sequencing coverage reduces the power to identify rare variants, affecting especially the heterozygosity (*H*) estimates. Dots show *H* and mean sequencing coverage for individuals of different populations, and the dashed lines the linear models fitted to each subset. For the Arctic samples, the correlation is significantly different from zero (p = 0.0001, Pearson’s product-moment correlation); for the Baltic, Ladoga and Saimaa samples, the correlation is weak or absent (p = 0.0271, p = 0.1570 and p = 0.8543, respectively).

**Fig. S3:**
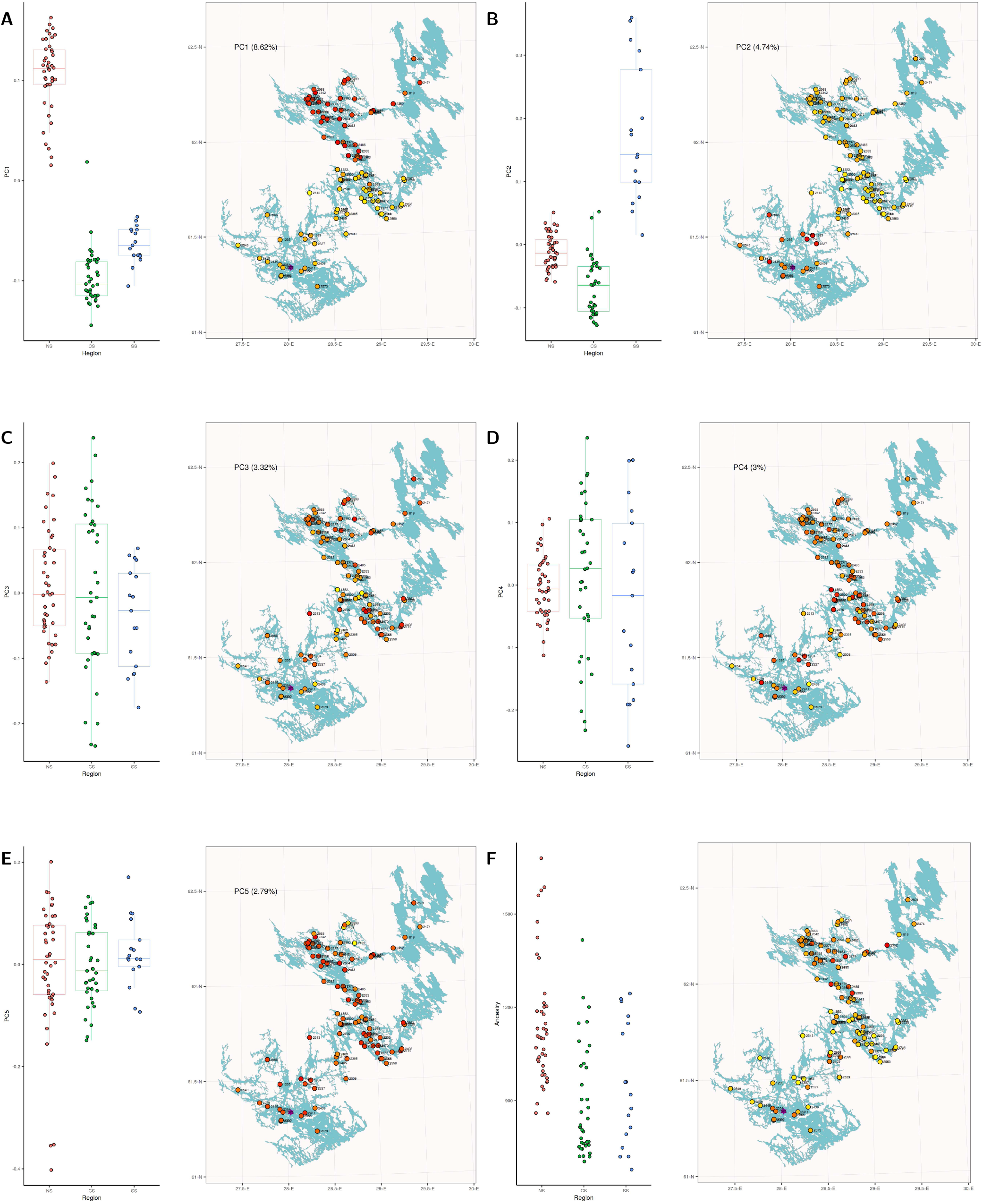
Distribution of PCs and coancestry on the map. (A-E) The distribution of the first five PCs in different subpopulations (left) and on the map (right). (F) The mean coancestry with the five NS samples identified as a diverged clade in the fineSTRUCTURE analysis. The dots on the map are colored by the PC/coancestry value, from yellow (low) to red (high).

**Fig. S4:**
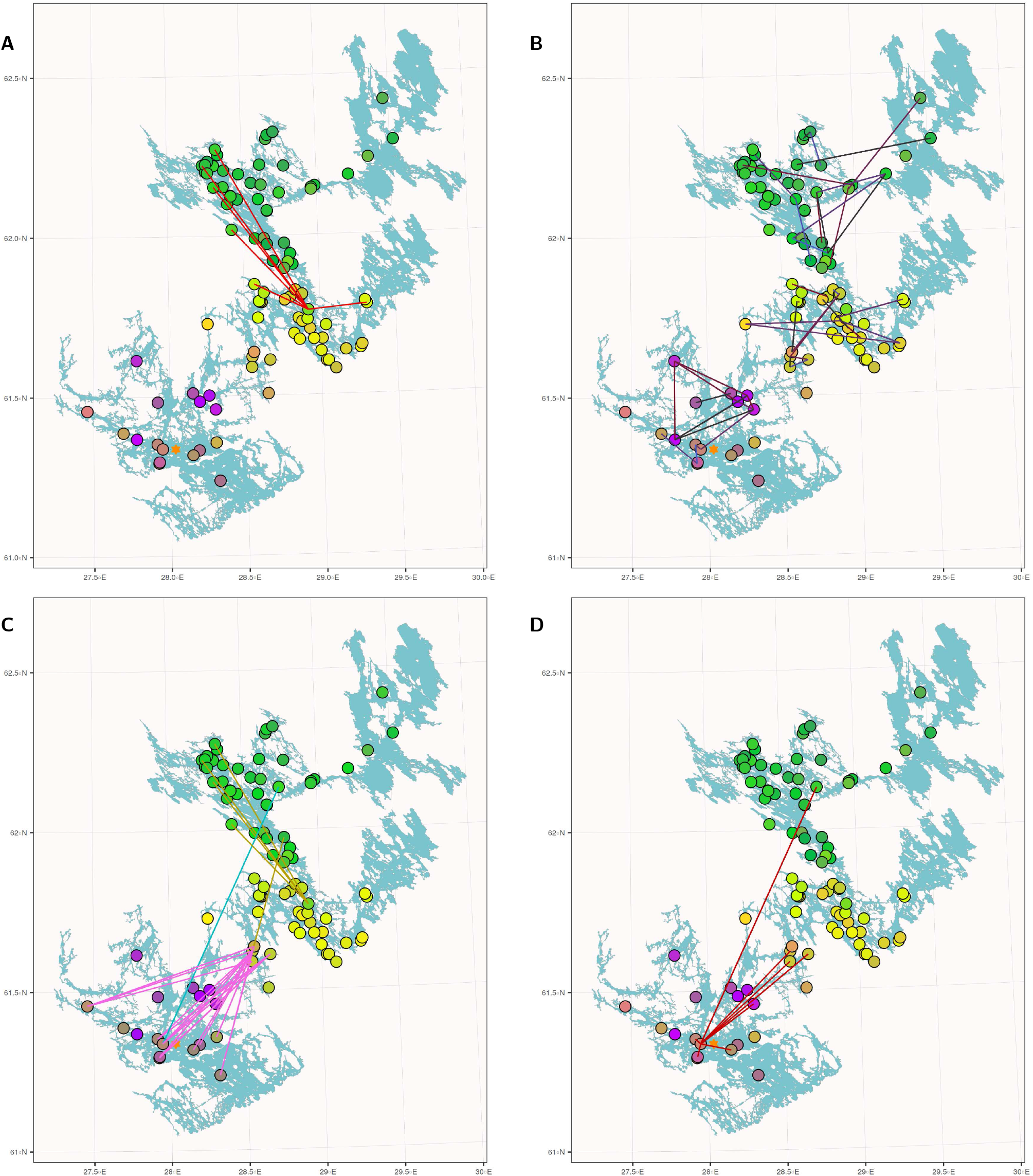
Samples with very high coancestry estimates. (A) Sample pairs of ‘2569’ with coancestry above 1500. (B) 25, 24 and 24 sample pairs, connected with lines, in the northern, central and southern part of Saimaa show coancestry above 2000. Line colors match the scale in Fig. 3D. (C) 29 sample pairs show coancestry above 1500 despite coming from different parts of the lake. Lines are colored by the subpopulations. (D) Sample pairs of ‘2575’ with coancestry above 1500.

**Fig. S5:**
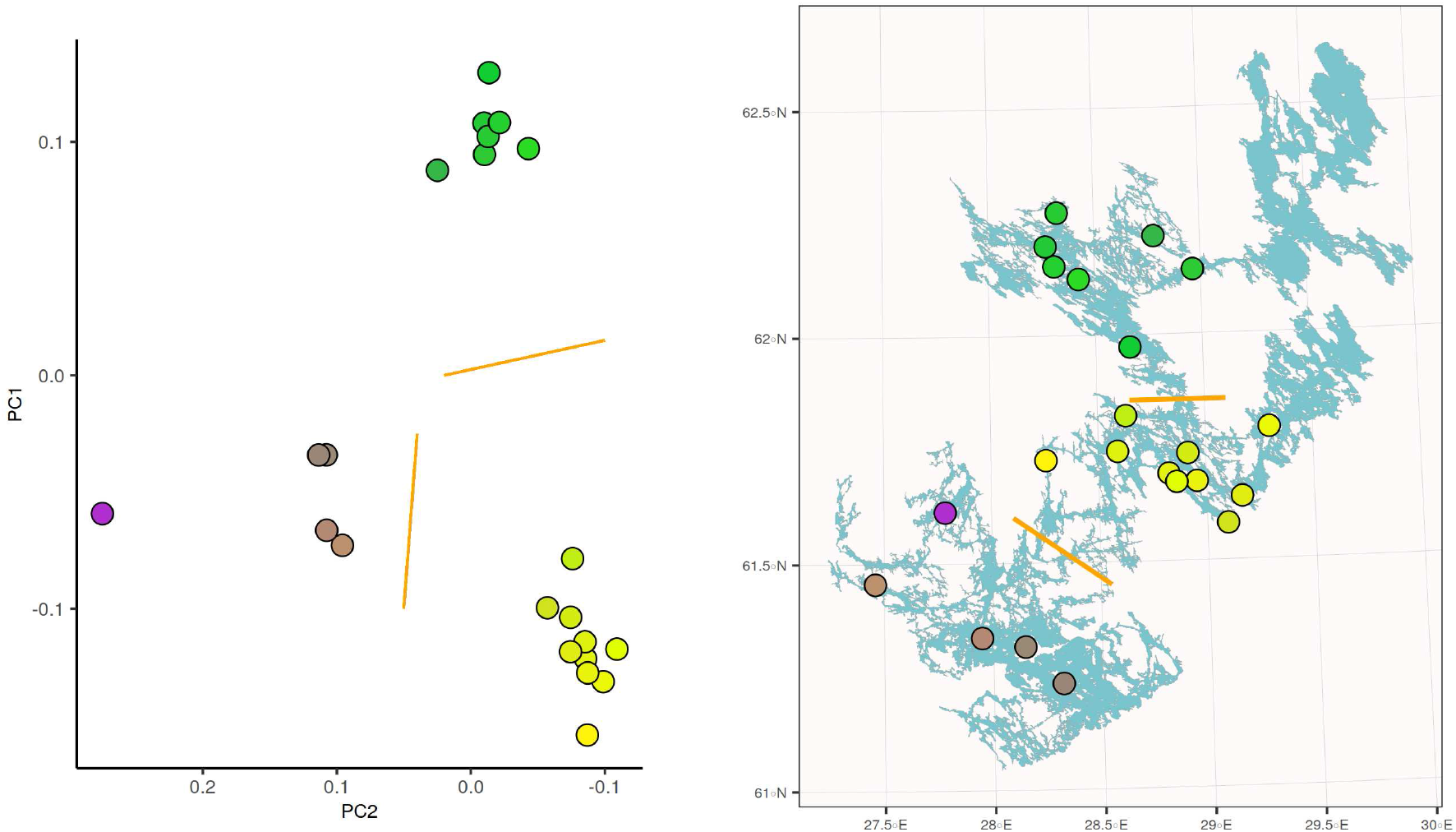
Saimaa seals born in the 2010s chosen to represent the current population.

**Fig. S6:**
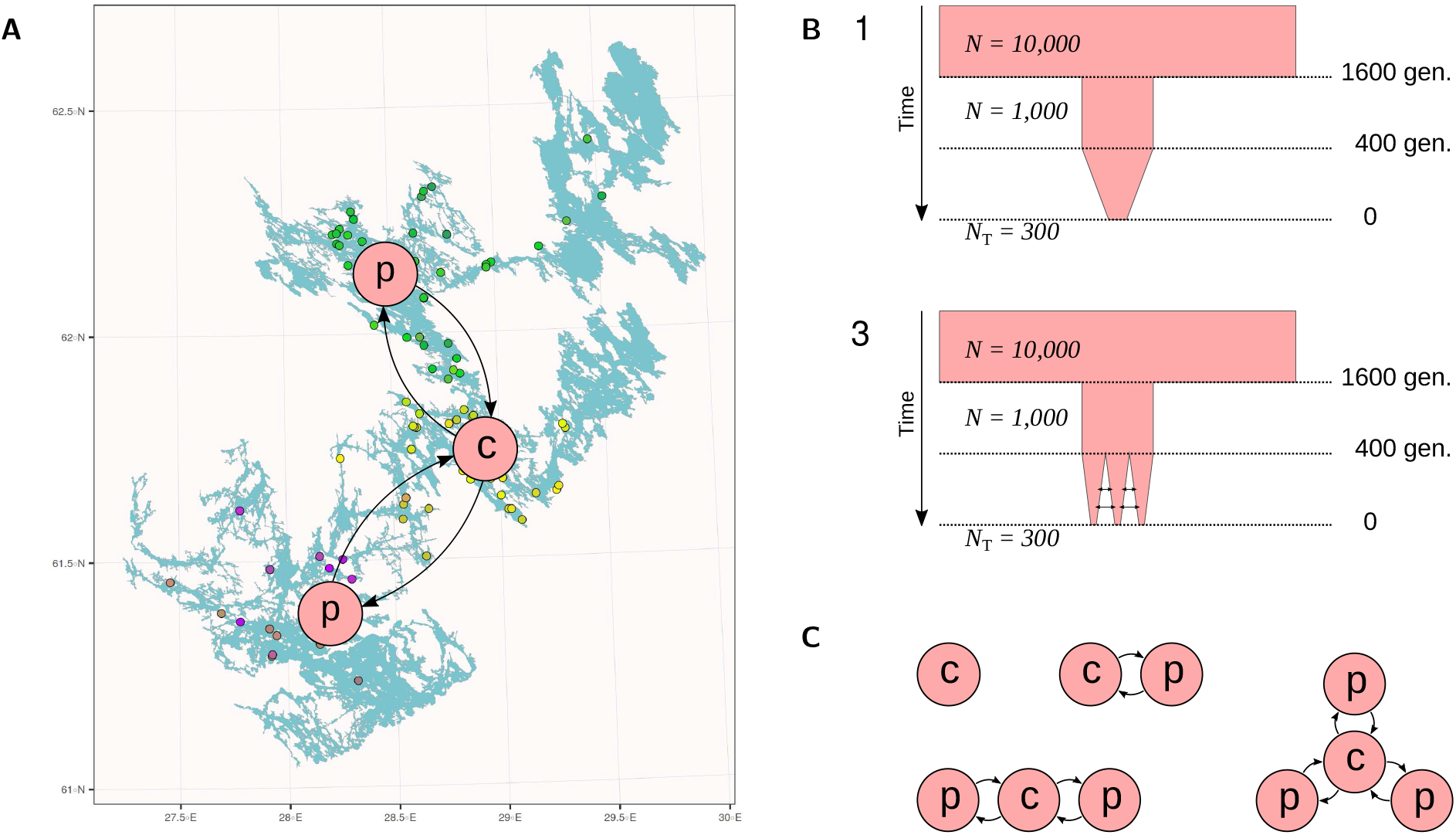
Simulation of substructured populations. (A) The Saimaa ringed seal population shows strong differentiation with limited historical migration between the subpopulations. Due to the fragmented shape of the lake, the central subpopulation (c) connects the two peripheric (p) parts of the lake. (B) We simulated synthetic data for a population divided into *N* =1..4 subpopulations that were either connected with migration (*m* > 0) or evolving in isolation (*m* = 0). The population was founded by an initial population of 1,000 individuals 1600 generations ago and this was then reduced to 300 individuals during the last 400 generations. (C) Schematic view of the star-shaped migration patterns. Peripheric (p) and central (c) subpopulations behave differently when the number of subpopulation is three or higher.

**Fig. S7:**
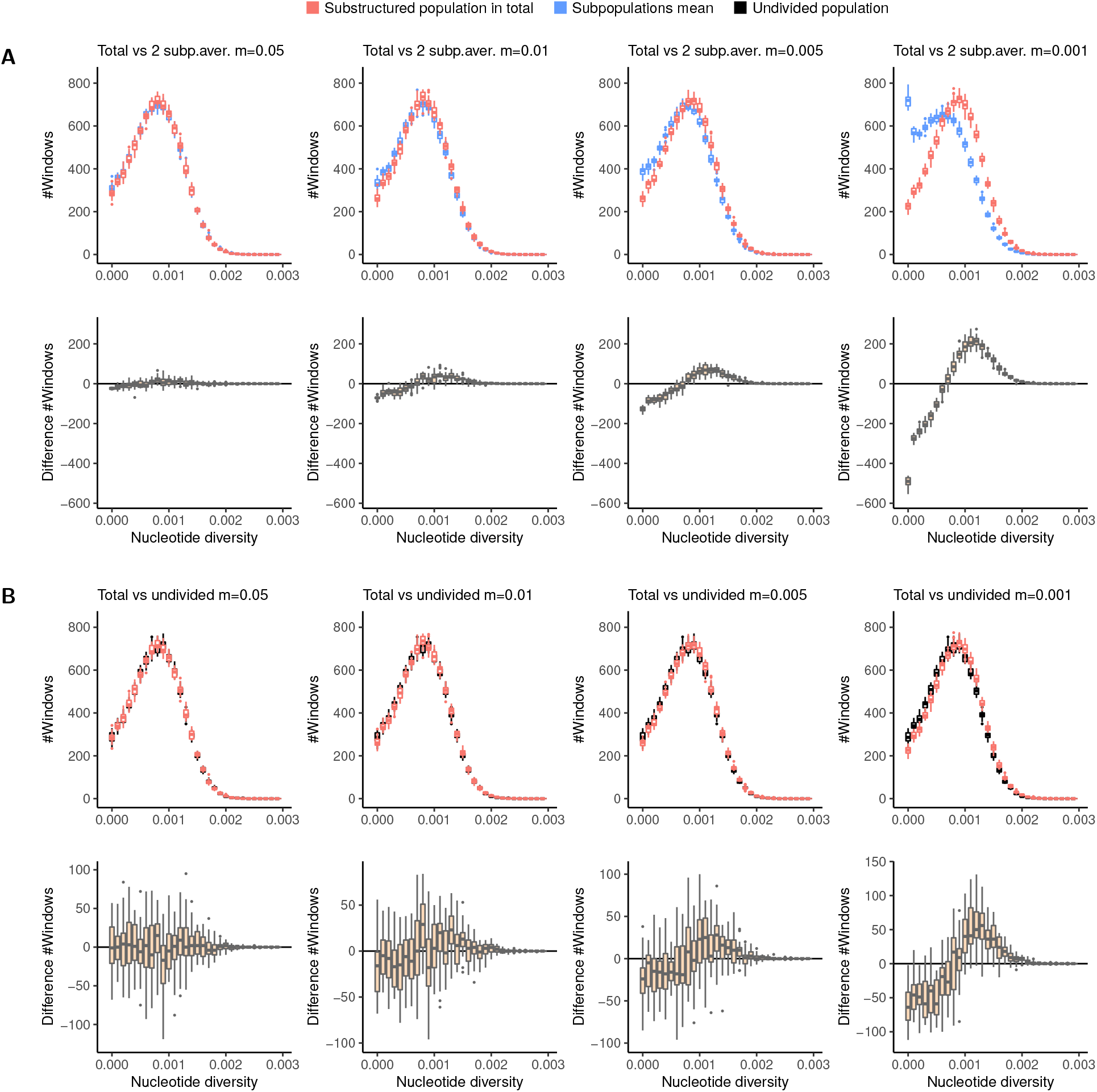
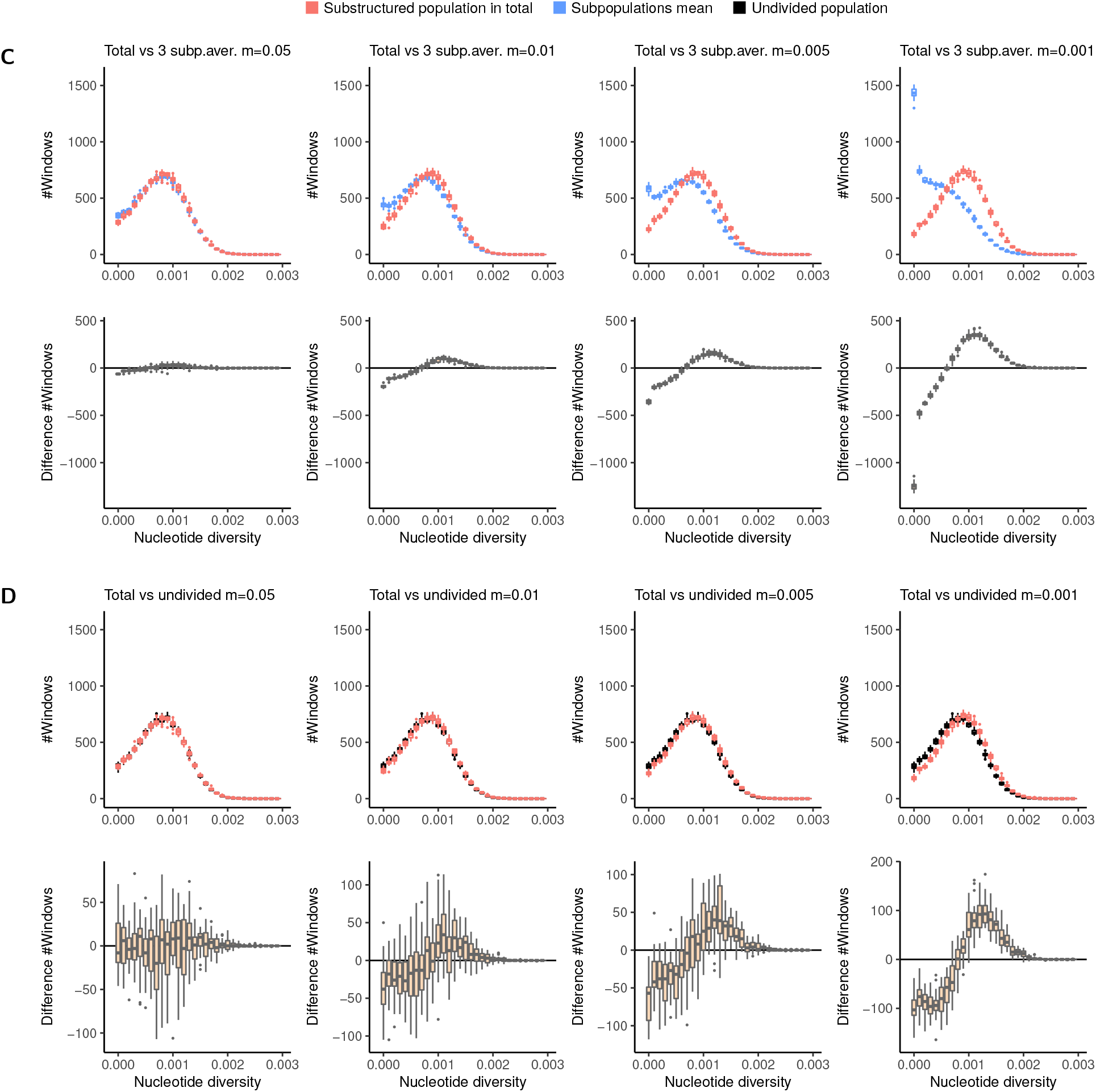
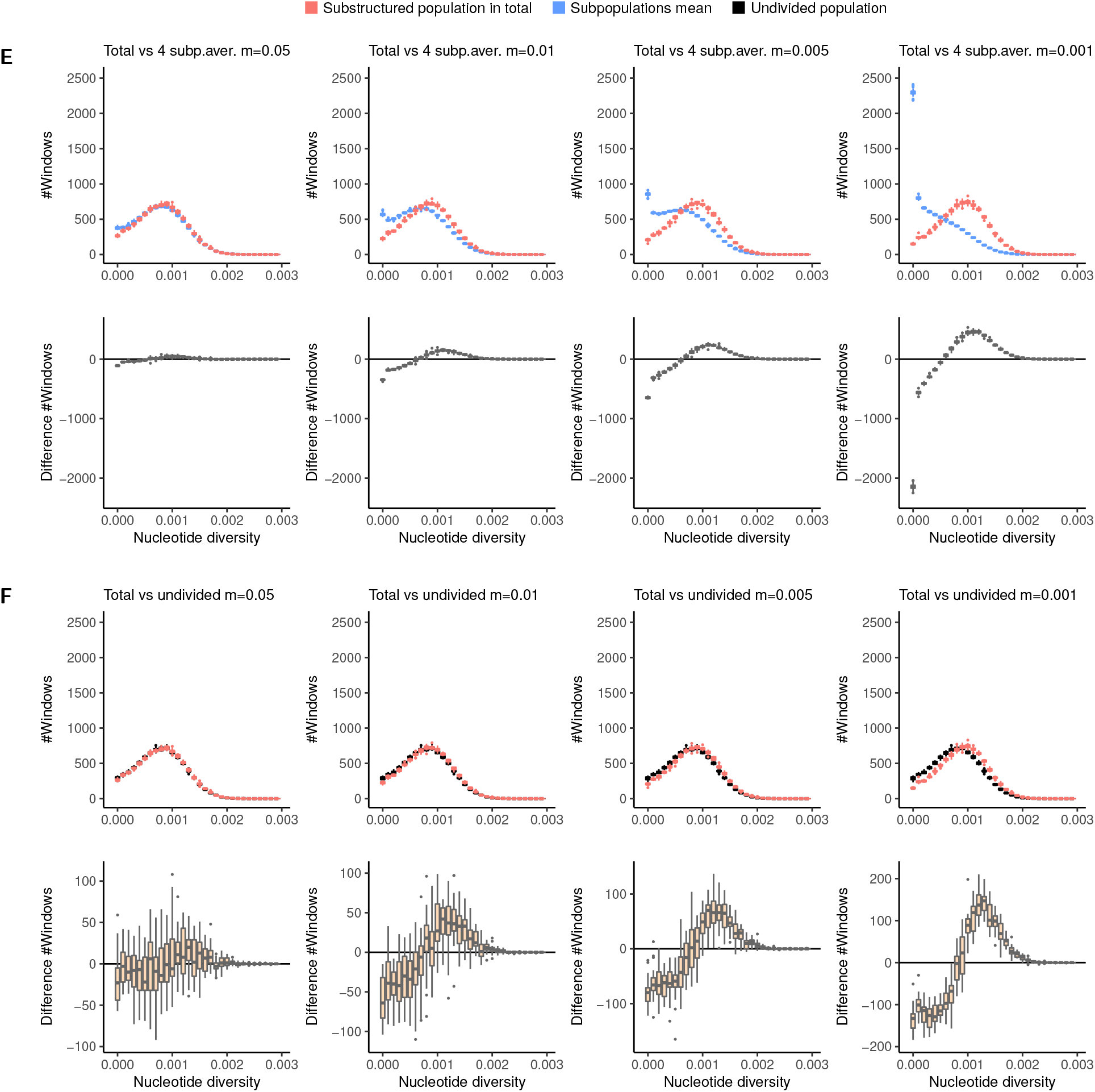
Preservation of variation in structured populations. (A) Distribution of nucleotide diversity (*π*) within 250kbp genomic blocks. Boxplots in the upper row show the number of windows (y axis) in each category of *π* (x axis) for the total population (red) and the average of two subpopulations. The panels (from left to right) show the results for migration rates 0.05, 0.01, 0.005 and 0.001, each consisting of 25 replicates. Boxplots in the lower row show the difference in number of windows between the two categories. Negative numbers indicate that the total population has fewer counts in that category than the subpopulations; positive numbers indicate increase in the total population. (B) Boxplots in the upper row show the number of windows in each category of *π* for the substructured population in total (red) and an undivided population (black). As an undivided population cannot contain migration, the same black distribution is repeated in different panels. Boxplots in the lower row show the difference in number of windows between the two categories. (C) Distribution of nucleotide diversity (*π*) within 250kbp genomic blocks when the population is divided into three subpopulations. (D) Comparison to an undivided population. (E) Distribution of nucleotide diversity (*π*) within 250kbp genomic blocks when the population is divided into four subpopulations. (F) Comparison to an undivided population.

**Table S1:** Sequencing coverage of the four-subspecies data (left) and the Saimaa data (right). The coverage was computed for the first contig (20.5 Mbp) from bam files, including all sites.

| Sample | Population | bam depth |  |  |  |  |
| --- | --- | --- | --- | --- | --- | --- |
| PH001 | Arctic/Ittoqqortoormiit | 3.442 |  |  |  |  |
| PH002 | Arctic/Ittoqqortoormiit | 2.643 |  |  |  |  |
| PH003 | Arctic/Ittoqqortoormiit | 4.254 |  |  |  |  |
| PH006 | Arctic/Ittoqqortoormiit | 3.759 |  |  |  |  |
| PH044 | Arctic/Ittoqqortoormiit | 5.888 |  |  |  |  |
| PH067 | Arctic/Qaanaaq | 6.703 |  |  |  |  |
| PH083 | Arctic/Ittoqqortoormiit | 5.110 |  |  |  |  |
| PH086 | Arctic/Ittoqqortoormiit | 5.514 |  |  |  |  |
| PH087 | Arctic/Ittoqqortoormiit | 5.230 |  |  |  |  |
| PH090 | Arctic/Ittoqqortoormiit | 5.183 |  |  |  |  |
| PH109 | Arctic/Qeqertarsuaq | 4.884 |  |  |  |  |
| PH112 | Arctic/Qeqertarsuaq | 6.248 |  |  |  |  |
| PH116 | Arctic/Qeqertarsuaq | 5.202 |  |  |  |  |
| PH130 | Arctic/Kangia | 5.975 |  |  |  |  |
| PH157 | Arctic/Holman | 5.316 |  |  |  |  |
| PH158 | Arctic/Holman | 4.912 |  |  |  |  |
| PH161 | Arctic/Holman | 5.088 |  |  |  |  |
| PH176 | Arctic/Ulukhaktok | 6.220 |  |  |  |  |
| PH185 | Arctic/Ulukhaktok | 5.523 |  |  |  |  |
| BG-01 | Baltic | 9.497 |  |  |  |  |
| BG-02 | Baltic | 8.727 |  |  |  |  |
| BG-03 | Baltic | 25.012 |  |  |  |  |
| BG-09 | Baltic | 11.576 |  |  |  |  |
| BG-10 | Baltic | 8.583 |  |  |  |  |
| BG-12 | Baltic | 10.462 |  |  |  |  |
| H2210 | Baltic | 19.518 |  |  |  |  |
| N160 | Baltic | 2.993 |  |  |  |  |
| N210 | Baltic | 10.508 |  |  |  |  |
| N3-09 | Baltic | 9.255 |  |  |  |  |
| N7-07 | Baltic | 8.377 |  |  |  |  |
| NN3-07 | Baltic | 11.388 |  |  |  |  |
| L1501 | Ladoga | 10.545 |  |  |  |  |
| L1683 | Ladoga | 12.949 |  |  |  |  |
| L1684 | Ladoga | 4.410 |  |  |  |  |
| LG-01 | Ladoga | 26.864 |  |  |  |  |
| LG-02 | Ladoga | 8.870 |  |  |  |  |
| LG-03 | Ladoga | 8.174 |  |  |  |  |
| LG-04 | Ladoga | 8.630 |  |  |  |  |
| LG-05 | Ladoga | 8.490 |  |  |  |  |
| LG-06 | Ladoga | 8.645 |  |  |  |  |
| 258 | Saimaa | 13.787 |  |  |  |  |
| 292 | Saimaa | 13.671 |  |  |  |  |
| 819 | Saimaa | 11.491 |  |  |  |  |
| 1090 | Saimaa | 8.663 |  |  |  |  |
| 1271 | Saimaa | 7.451 |  |  |  |  |
| 1482 | Saimaa | 13.241 |  |  |  |  |
| 1983 | Saimaa | 22.659 |  |  |  |  |
| 2259 | Saimaa | 6.670 |  |  |  |  |
| 2343 | Saimaa | 8.569 |  |  |  |  |
| 2357 | Saimaa | 8.169 |  |  |  |  |
| 2399 | Saimaa | 11.892 |  |  |  |  |
| 2436 | Saimaa | 8.646 |  |  |  |  |
| 2449 | Saimaa | 12.885 |  |  |  |  |
| 2474 | Saimaa | 11.430 |  |  |  |  |
| 2496 | Saimaa | 8.309 |  |  |  |  |
| 2528 | Saimaa | 13.333 |  |  |  |  |
| 2564 | Saimaa | 8.831 |  |  |  |  |
| 2585 | Saimaa | 10.677 |  |  |  |  |
| 2614 | Saimaa | 12.772 |  |  |  |  |
|  |  |  | Sample | bam depth | Sample | bam depth |
|  |  |  | 297 | 3.643 | 2405 | 1.798 |
|  |  |  | 298 | 6.986 | 2411 | 4.682 |
|  |  |  | 1235 | 4.652 | 2421 | 2.225 |
|  |  |  | 1262 | 6.206 | 2422 | 5.195 |
|  |  |  | 1270 | 6.638 | 2426 | 4.626 |
|  |  |  | 1488 | 9.300 | 2427 | 10.610 |
|  |  |  | 1552 | 8.174 | 2432 | 7.058 |
|  |  |  | 1956 | 3.269 | 2433 | 5.591 |
|  |  |  | 1957 | 7.442 | 2440 | 6.053 |
|  |  |  | 1977 | 1.260 | 2445 | 4.460 |
|  |  |  | 2119 | 2.696 | 2452 | 2.797 |
|  |  |  | 2221 | 1.209 | 2454 | 6.967 |
|  |  |  | 2232 | 8.215 | 2456 | 2.619 |
|  |  |  | 2233 | 7.270 | 2472 | 5.434 |
|  |  |  | 2277 | 6.831 | 2477 | 3.938 |
|  |  |  | 2327 | 1.113 | 2485 | 1.176 |
|  |  |  | 2332 | 2.466 | 2497 | 3.276 |
|  |  |  | 2333 | 0.434 | 2499 | 5.857 |
|  |  |  | 2337 | 5.460 | 2500 | 4.045 |
|  |  |  | 2340 | 7.151 | 2501 | 6.541 |
|  |  |  | 2341 | 4.643 | 2504 | 3.594 |
|  |  |  | 2342 | 7.152 | 2506 | 8.741 |
|  |  |  | 2348 | 5.958 | 2509 | 5.502 |
|  |  |  | 2350 | 6.187 | 2511 | 5.242 |
|  |  |  | 2352 | 0.836 | 2513 | 6.235 |
|  |  |  | 2353 | 2.023 | 2514 | 5.089 |
|  |  |  | 2354 | 1.210 | 2527 | 2.055 |
|  |  |  | 2356 | 5.142 | 2549 | 4.784 |
|  |  |  | 2363 | 4.621 | 2550 | 7.326 |
|  |  |  | 2375 | 6.257 | 2551 | 6.412 |
|  |  |  | 2376 | 3.183 | 2555 | 1.772 |
|  |  |  | 2377 | 0.558 | 2557 | 5.416 |
|  |  |  | 2379 | 8.209 | 2563 | 5.927 |
|  |  |  | 2380 | 5.550 | 2565 | 8.138 |
|  |  |  | 2382 | 7.823 | 2566 | 8.144 |
|  |  |  | 2383 | 5.900 | 2567 | 6.741 |
|  |  |  | 2392 | 8.311 | 2568 | 6.452 |
|  |  |  | 2393 | 7.478 | 2569 | 9.112 |
|  |  |  | 2395 | 1.123 | 2570 | 8.488 |
|  |  |  | 2396 | 5.503 | 2572 | 5.062 |
|  |  |  | 2397 | 8.362 | 2573 | 4.459 |
|  |  |  | 2402 | 3.998 | 2574 | 8.231 |
|  |  |  | 2404 | 4.262 | 2575 | 0.342 |

**Table S2:** For each sample, sampling region and year, age at death, and heterozygosity across the longest 100 contigs.

| Sample | Region | Year | Age | $H$ | Sample | Region | Year | Age | $H$ |
| --- | --- | --- | --- | --- | --- | --- | --- | --- | --- |
| 819 | NS | 1985 | 0 | 0.001207 | 2411 | SS | 2006 | 0 | 0.001323 |
| 1090 | CS | 1986 | 0 | 0.001541 | 2421 | CS | 2006 | 16 | 0.001376 |
| 1235 | SS | 1987 | 0 | 0.001337 | 2422 | NS | 2007 | 0 | 0.001338 |
| 1262 | NS | 1988 | 0 | 0.001373 | 2426 | NS | 2007 | 0 | 0.001319 |
| 1482 | NS | 1994 | 0 | 0.001322 | 2427 | CS | 2007 | 0 | 0.001365 |
| 1488 | SS | 1994 | 21 | 0.001412 | 2432 | NS | 2007 | 0 | 0.001298 |
| 1552 | CS | 1995 | 0 | 0.001207 | 2433 | NS | 2007 | 0 | 0.001456 |
| 1956 | NS | 1997 | 0 | 0.001383 | 2436 | SS | 2007 | 0 | 0.001139 |
| 1957 | SS | 1997 | 0 | 0.001084 | 2440 | NS | 2007 | 0 | 0.001350 |
| 1977 | CS | 1997 | 0 | 0.001341 | 2445 | CS | 2007 | 0 | 0.001364 |
| 1983 | NS | 1997 | 0 | 0.001047 | 2449 | SS | 2007 | 6 | 0.001084 |
| 2119 | CS | 1998 | 6 | 0.001070 | 2452 | NS | 2008 | 0 | 0.001134 |
| 2221 | NS | 2000 | 28 | 0.001379 | 2454 | NS | 2008 | 0 | 0.001297 |
| 2232 | NS | 2000 | 5 | 0.001375 | 2456 | CS | 2008 | 0 | 0.001330 |
| 2233 | CS | 2000 | 0 | 0.001578 | 2472 | CS | 2008 | 0 | 0.001192 |
| 2259 | SS | 2001 | 0 | 0.001429 | 2474 | NS | 2009 | 2 | 0.001480 |
| 2277 | NS | 2001 | NA | 0.001508 | 2477 | CS | 2009 | 0 | 0.001256 |
| 2327 | SS | 2002 | 0 | 0.001241 | 2485 | NS | 2009 | 5 | 0.001438 |
| 2332 | SS | 2003 | 1 | 0.001563 | 2496 | CS | 2009 | 0 | 0.001142 |
| 2333 | NS | 2003 | NA | 0.001378 | 2497 | CS | 2010 | 3 | 0.001305 |
| 2337 | NS | 2003 | 12 | 0.001373 | 2499 | NS | 2010 | 0 | 0.001119 |
| 2340 | NS | 2003 | 0 | 0.001234 | 2500 | CS | 2010 | 0 | 0.001309 |
| 2341 | NS | 2003 | 0 | 0.001175 | 2501 | NS | 2010 | 6 | 0.001417 |
| 2342 | NS | 2003 | 0 | 0.001227 | 2504 | NS | 2010 | 33 | 0.001305 |
| 2343 | CS | 2003 | 0 | 0.001352 | 2506 | NS | 2010 | 0 | 0.001424 |
| 2348 | NS | 2003 | 1 | 0.001378 | 2509 | CS | 2010 | 0 | 0.001311 |
| 2350 | NS | 2003 | 0 | 0.001393 | 2511 | NS | 2010 | 0 | 0.001244 |
| 2352 | NS | 2003 | 0 | 0.001435 | 2513 | CS | 2010 | 0 | 0.001191 |
| 2353 | SS | 2004 | 1 | 0.001507 | 2514 | CS | 2011 | 20 | 0.001244 |
| 2354 | SS | 2004 | 1 | 0.001529 | 2527 | CS | 2011 | 0 | 0.001201 |
| 2356 | NS | 2004 | 0 | 0.001360 | 2528 | CS | 2011 | 2 | 0.001331 |
| 2357 | SS | 2004 | 1 | 0.001083 | 2549 | SS | 2012 | 0 | 0.001081 |
| 2363 | NS | 2004 | 0 | 0.001234 | 2550 | CS | 2012 | 0 | 0.001394 |
| 2375 | CS | 2005 | 0 | 0.001302 | 2551 | NS | 2012 | 0 | 0.001073 |
| 2376 | CS | 2005 | 0 | 0.001351 | 2555 | SS | 2012 | 0 | 0.001276 |
| 2377 | NS | 2005 | 0 | 0.001433 | 2557 | NS | 2012 | 0 | 0.001406 |
| 2379 | NS | 2005 | 0 | 0.001526 | 2563 | CS | 2013 | NA | 0.001046 |
| 2380 | NS | 2005 | 0 | 0.000997 | 2564 | CS | 2013 | NA | 0.001403 |
| 2382 | NS | 2005 | 0 | 0.001415 | 2565 | CS | 2013 | NA | 0.001232 |
| 2383 | CS | 2005 | NA | 0.001410 | 2566 | SS | 2013 | 0 | 0.001305 |
| 2392 | NS | 2005 | 0 | 0.001301 | 2567 | CS | 2013 | 0 | 0.001345 |
| 2393 | NS | 2005 | 0 | 0.001483 | 2568 | NS | 2013 | 0 | 0.001479 |
| 2395 | CS | 2006 | 1 | 0.001477 | **2569** | CS | 2013 | 0 | **0.001667** |
| 2396 | NS | 2006 | 5 | 0.001114 | 2570 | CS | 2013 | 0 | 0.001357 |
| 2397 | CS | 2006 | 0 | 0.001210 | 2572 | CS | 2013 | 0 | 0.001226 |
| 2399 | NS | 2006 | 0 | 0.001128 | 2573 | SS | 2013 | 0 | 0.001193 |
| 2402 | CS | 2006 | 0 | 0.001023 | 2574 | CS | 2013 | 0 | 0.001375 |
| 2404 | CS | 2006 | 0 | 0.001289 | 2575 | SS | 2013 | 0 | 0.001536 |
| 2405 | CS | 2006 | 0 | 0.001276 | 2585 | NS | 2013 | 0 | 0.001432 |

## References

1. Frankham, R. Effective population size/adult population size ratios in wildlife: a review. Genet. Res. 66, 95–107 (1995).

2. Palstra, F. P. & Ruzzante, D. E. Genetic estimates of contemporary effective population size: what can they tell us about the importance of genetic stochasticity for wild population persistence? Mol. Ecol. 17, 3428–3447 (2008).

3. Hare, M. P. et al. Understanding and estimating effective population size for practical application in marine species management. Conserv. Biol. 25, 438–449 (2011).

4. Wright, S. Evolution in mendelian populations. Genetics 16, 97–159 (1931).

5. Fisher, R. A. The distribution of gene ratios for rare mutations. Proceedings of the Royal Society of Edinburgh 50, 204–219 (1931).

6. Waples, R. S. Spatial-temporal stratifications in natural populations and how they affect understanding and estimation of effective population size. Mol. Ecol. Resour. 10, 785–796 (2010).

7. Wright, S. Size of population and breeding structure in relation to evolution. Science 87, 430–431 (1938).

8. Wakeley, J. Coalescent Theory: An Introduction (W. H. Freeman, 2008), 1 edn.

9. NAMMCO. Ringed seals. https://nammco.no/topics/ringed-seal.

10. Kunnasranta, M. et al. Sealed in a lake — biology and conservation of the endangered Saimaa ringed seal: A review. Biol. Conserv. 253, 108908 (2021).

11. Valtonen, M., Palo, J. U., Ruokonen, M., Kunnasranta, M. & Nyman, T. Spatial and temporal variation in genetic diversity of an endangered freshwater seal. Conserv. Genet. 13, 1231–1245 (2012).

12. Valtonen, M. et al. Causes and consequences of fine-scale population structure in a critically endangered freshwater seal. BMC Ecol. 14, 22 (2014).

13. Li, H. SNPable. http://lh3lh3.users.sourceforge.net/snpable.shtml. Accessed: 2021-10-20.

14. Smit, A., Hubley, R. & Green, P. RepeatMasker open-3.0 (1996).

15. Li, H. Aligning sequence reads, clone sequences and assembly contigs with bwa-mem (2013). http://arxiv.org/abs/1303.3997.

16. Danecek, P. et al. Twelve years of SAMtools and BCFtools. Gigascience 10, giab008 (2021).

17. Korneliussen, T., Albrechtsen, A. & Nielsen, R. ANGSD: Analysis of next generation sequencing data. BMC Bioinformatics 15, 356 (2014).

18. Browning, B. L., Tian, X., Zhou, Y. & Browning, S. R. Fast two-stage phasing of large-scale sequence data. Am. J. Hum. Genet. 108, 1880–1890 (2021).

19. Schiffels, S. & Wang, K. MSMC and MSMC2: The multiple sequentially Markovian coalescent. Methods Mol. Biol. 2090, 147–166 (2020).

20. Liu, S. et al. Population genomics reveal recent speciation and rapid evolutionary adaptation in polar bears. Cell 157, 785–794 (2014).

21. Palo, J. U., Mäkinen, H. S., Helle, E., Stenman, O. & Väinölä, R. Microsatellite variation in ringed seals (Phoca hispida): genetic structure and history of the Baltic Sea population. Heredity 86, 609–617 (2001).

22. Koskinen, M. T. et al. Genetic lineages and postglacial colonization of grayling (thymallus thymallus, salmonidae) in europe, as revealed by mitochondrial DNA analyses. Mol. Ecol. 9, 1609–1624 (2000).

23. Nilsson, J. et al. Matrilinear phylogeography of atlantic salmon (salmo salar l.) in europe and postglacial colonization of the baltic sea area. Mol. Ecol. 10, 89–102 (2001).

24. Säisä, M. et al. Population genetic structure and postglacial colonization of Atlantic salmon (Salmo salar) in the Baltic Sea area based on microsatellite DNA variation. Can. J. Fish. Aquat. Sci. 62, 1887–1904 (2005).

25. Heino, M. T. et al. Museum specimens of a landlocked pinniped reveal recent loss of genetic diversity and unexpected population connections (2022).

26. Ceballos, F. C., Joshi, P. K., Clark, D. W., Ramsay, M. & Wilson, J. F. Runs of homozygosity: windows into population history and trait architecture. Nat. Rev. Genet. 19, 220–234 (2018).

27. Lawson, D. J., Hellenthal, G., Myers, S. & Falush, D. Inference of population structure using dense haplotype data. PLoS Genet. 8, e1002453 (2012).

28. Bhatia, G., Patterson, N., Sankararaman, S. & Price, L. Estimating and interpreting FST: the impact of rare variants. Genome Res. 23, 1514–1521 (2013).

29. Koskela, J. T., Kunnasranta, M., Hämäläinen, E. & Hyvärinen, H. Movements and use of haul-out sites of radio-tagged Saimaa ringed seal (Phoca hispida saimensis Nordq.) during the open-water season. Ann. Zool. Fennici 39, 59–67 (2002).

30. Wintle, B. A. et al. Global synthesis of conservation studies reveals the importance of small habitat patches for biodiversity. Proc. Natl. Acad. Sci. U. S. A. 116, 909–914 (2019).

31. Nyman, T. et al. Demographic histories and genetic diversities of Fennoscandian marine and landlocked ringed seal subspecies. Ecol. Evol. 4, 3420–3434 (2014).

32. Kovacs, K. M. et al. Global threats to pinnipeds. Mar. Mamm. Sci. 28, 414–436 (2012).

33. Chin, C.-S. et al. Phased diploid genome assembly with single-molecule real-time sequencing. Nat. Methods 13, 1050–1054 (2016).

34. Walker, B. J. et al. Pilon: an integrated tool for comprehensive microbial variant detection and genome assembly improvement. PLoS One 9, e112963 (2014).

35. McKenna, A. et al. The genome analysis toolkit: a MapReduce framework for analyzing next-generation DNA sequencing data. Genome Res. 20, 1297–1303 (2010).

36. Browning, B. L. & Browning, S. R. Genotype imputation with millions of reference samples. Am. J. Hum. Genet. 98, 116–126 (2016).

37. Narasimhan, V. et al. BCFtools/RoH: a hidden Markov model approach for detecting autozygosity from next-generation sequencing data. Bioinformatics 32, 1749–1751 (2016).

38. Lawrence, M. et al. Software for computing and annotating genomic ranges. PLoS Comput. Biol. 9, e1003118 (2013).

39. Danecek, P. et al. The variant call format and VCFtools. Bioinformatics 27, 2156–2158 (2011).

40. Patterson, N., Price, A. L. & Reich, D. Population structure and eigenanalysis. PLoS Genet. 2, e190 (2006).

41. Kelleher, J., Etheridge, A. M. & McVean, G. Efficient coalescent simulation and genealogical analysis for large sample sizes. PLoS Comput. Biol. 12, e1004842 (2016).

